# Retinal scRNAseq reveals cell-type-specific responses to bacterial infection

**DOI:** 10.64898/2026.05.26.725952

**Authors:** Sierra Foshe, Helen Rossmiller, Jacob K Sterling, Ellen White, Michelle C Callegan, Yuyan Cheng, Elizabeth Grice, Joshua L Dunaief

## Abstract

**Purpose:** Endophthalmitis is a serious complication of intraocular surgery due to the risk of irreversible retinal damage. Because retinal cell populations are highly heterogenous, single-cell resolution is required to uncover the detailed mechanisms of infection response. Using a mouse model of bacterial endophthalmitis, we investigated transcriptional changes across resident and infiltrating cell types in the retina.

**Methods:** A methicillin-sensitive strain of *Staphylococcus aureus* was isolated from a patient with endophthalmitis. Adult C57Bl/6J mice received an intravitreal injection of phosphate-buffer solution (PBS) with or without 5000 CFU *S. aureus* (n=3 per group). 24 hours later, retinas were isolated and single-cell suspensions were sent to the Penn Genomic Core for sequencing with an Illumina NovaSeq 6000. After standard pre-processing of the data, differential genes and pathways were identified for each cell type (adjusted p < 0.01, log2FC > 1 or < -1).

**Results:** Our analysis identified all expected retinal cell types, including a population of infiltrating neutrophils in the infected retinas. We surveyed genes known to be upregulated at the bulk-retina level in this model (e.g. *Tlr2, Nlrp3, Il1b*), and found that infiltrating cells mainly drove this expression. Several genes were altered across nearly all retinal cell types, including upregulation of *Hsph1* and *Stat3*. Muller glia downregulated *Gpx4* while upregulating *Acsl4* and iron importers *Tfrc, Zip14*, and *Dmt1*. Top pathways for macrophages/microglia included chemotaxis, cell-cell adhesion, and wound healing. Vascular cells upregulated angiogenesis-related genes. Cellular respiration was a commonly affected pathway across several neuronal populations, with most genes decreasing.

**Conclusions:** This study advances our understanding of the pathobiology of bacterial endophthalmitis. Muller glia appear to be undergoing ferroptosis, potentially while activating a program to sequester iron away from bacteria. Decreased cellular respiration may indicate hypoxia among neurons. Our results reveal several trends in the retinal response to infection, including iron dysregulation and hypoxia. Understanding these cell-type-specific responses to endophthalmitis may help design therapies to combine with antibiotics.

## Introduction

Infection is one of the most serious complications of surgical procedures. Bacterial endophthalmitis, infection within the eye, is a medical emergency. Retinal damage is difficult to prevent: Infection can progress rapidly and irreversibly, as the central nervous system is non-regenerative. Endophthalmitis is primarily caused by intraocular operations like cataract removal or intravitreal drug injections, but can also result from traumatic injury or systemic infection spreading to the eye.^1^

When a patient presents with suspected endophthalmitis, a sample of the vitreous fluid is taken for culture and antibiotic treatment is typically started before receiving results. It takes several days to identify the causative pathogen and its antibiotic susceptibility, potentially delaying effective treatment. The most severe cases of endophthalmitis are resolved by enucleation, removal of the eye. Even with less severe cases, up to a third of patients experience significant, permanent vision loss.^2^

The virulent pathogen *Staphylococcus aureus* can cause severe endophthalmitis infection.^3^ It is the causative species in 10% of post-cataract cases and 25% of endogenous (arising from systemic infection) cases.^1^ *S. aureus* is notorious for developing antibiotic resistance, making it particularly difficult to treat and leading to worse outcomes. Up to 40% of *S. aureus* isolates from endophthalmitis patients are the highly pathogenic methicillin-resistant strain (MRSA).^2^ Full resistance to vancomycin, the “last resort” antibiotic for resistant infections, has recently appeared in *S. aureus* and other species.^2^ The combination of high virulence, increasing resistance, and substantial disease burden make *S. aureus* of particular importance in endophthalmitis research.

A well-characterized mouse model of *S. aureus* endophthalmitis has been used to study disease progression. Within 24h of infection, bacterial load increases over 500-fold and overt signs are present: the cornea is cloudy and white, and retinal lamination is disrupted.^4–6^ Pro-inflammatory cytokines including IL-1β and IL-6 are upregulated, myeloid cells infiltrate the retina, and retinal function is almost completely abolished.^4^

A combination of bacterial toxins and runaway inflammation quickly leads to retinal detachment and degeneration. *S. aureus* produces a variety of virulence factors including Panton-Valentine leukocidin, alpha-toxin, and Protein A, which are each sufficient to induce apoptosis and retinal dysfunction.^7,8^

Bacterial growth inside the eye can also indirectly lead to retinal degeneration by eliciting a rampant inflammatory response.^9,10^ The recruitment of immune cells to clear bacteria can inadvertently damage host cells. Visual outcome in murine models is improved by reducing inflammation through knock-out of the chemokine CXCL1,^11^ or immune cell receptor TLR4.^12^ Retinal degeneration therefore results from both direct bacterial toxicity and secondary inflammatory injury.

However, some amount of inflammation is necessary for responding to infection. Outcomes can conversely be worsened by knock-out of key immune regulators like TNF-α and IL-1β.^13,14^ Therefore, instead of targeting a central mediator of immune response, blocking a pathway that contributes to the toxic cycle of inflammation may be more effective.

The goal of this study was to investigate how retinal cells respond to *S. aureus* infection, to identify novel targets for therapies.

## Methods

### Animals

Ten week-old C57BL/6J mice (strain number 000664) were purchased from Jackson Laboratory (Bar Harbor, ME) and housed in an Animal Biosafety Level-2 facility with *ad libitum* access to food and water on a 12h/12h light/dark schedule. A mixture of males and females were used in each group. All experiments were approved by the Institutional Animal Care and Use Committee (IACUC) of the University of Pennsylvania.

### Induction of endophthalmitis

Mice received 5000 CFU of *Staphylococcus aureus*, or PBS control, by intravitreal injection. *S. aureus* was a methicillin-sensitive strain derived from a patient with endophthalmitis.

Before injection, mice were anesthetized by intraperitoneal injection of ketamine/xylazine. Their pupils were dilated with tropicamide/phenylephrine hydrochloride. While proptosing the eye with curved forceps, 1 µL was injected into the vitreous cavity with a 32G needle microsyringe (Hamilton, Reno, NV). To protect the cornea while the mouse recovered from anesthesia, their eyes were covered with a layer of Lubricant PM Ointment (AACE Pharmaceuticals 71406-124-35).

### Immunohistochemistry (IHC)

At 24 hr post-injection, eyes were enucleated and fixed in 4% paraformaldehyde for 3 days at 4 °C. After washing with PBS, they were incubated in 30% sucrose overnight at 4 °C, then flash-frozen in Tissue-Tek OCT Compound and stored at -80 °C.

Tissue was cut into 10 µm sections with a Leica CM3050 cryostat. Slides were dried overnight and stored at -20 °C. After washing in PBS, slides were coverslipped with DAPI Fluoromount-G Mounting Medium (SouthernBiotech 0100-20) and imaged with a Nikon Eclipse 80i microscope.

### Single cell RNA sequencing (scRNAseq)

At 24 hours post-injection, eyes were enucleated with curved forceps and retinas were dissected and processed following an optimized protocol.^15^ Live/dead staining was performed with the LIVE/DEAD Cell Imaging Kit (ThermoFisher R37601) according to manufacturer’s directions. Single-cell suspensions were submitted to the Penn Genomic Core for sequencing on an Illumina NovaSeq 6000, using 10x Genomics Single Cell 3’ v3 chemistry.

### scRNAseq analysis

Reads were aligned with CellRanger v7.2.0. To remove ambient RNA background, data was pre-processed with CellBender v0.2.0^16^ before analysis with the R Seurat package. Further quality control was performed to remove doublets and cells with high mitochondrial gene expression. All six samples were integrated with IntegrateData. Cell type labeling was performed with the Azimuth package based on reference data from the Mouse Retina Cell Atlas (MRCA)^17^ and a dataset of non-neuronal cells after optic nerve injury.^18^ AggreagteExpression and FindMarkers (with DESeq2) were used to perform differential gene expression analysis. Differentially expressed genes were defined as having an adjusted p-value < 0.01 and log_2_ fold-change > 1 or < -1.

### MIO-M1 culture

The human immortalized Muller glia cell line MIO-M1 was originally obtained from University College, London, UK. Cells were cultured in 24-well plates with D10 media (DMEM + GlutaMax (Invitrogen, CA, USA), 10% fetal bovine serum (Hyclone, UT, USA), and penicillin/streptomycin) in a 37 °C, 5% CO_2_ incubator. Cells were incubated with or without 10 ng/mL active human IL-1β protein (Abcam ab259387) for 24 hours.

### Quantitative PCR (qPCR)

Cell lysates were collected and RNA isolation, reverse transcription, and qPCR were performed as previously described.^19^ Gene expression was measured using Taqman probes (Applied Biosystems, Foster City, CA): Euk 18s rRNA (endogenous control, Hs99999901_s1), *Il6* (interleukin-6, Hs00985639_m1), *Fpn* (ferroportin, Hs0020588_m1), *Slc39a14* (Zrt- and Irt-like protein 14 (*Zip14*), Hs00299262_m1), *Slc11a2* (divalent metal transport 1 (*Dmt1*), Hs00167206_m1), *Tfrc* (transferrin receptor, Hs00174609_m1), *Gpx4* (glutathione peroxidase 4, Hs00989766_g1) and *Acsl4* (acyl-CoA synthetase long chain family member 4, Hs00244871_m1).

## Results

### S. aureus endophthalmitis induces rampant retinal inflammation and neutrophil infiltration

Adult wild-type mice received an intravitreal injection of *S. aureus* or PBS (n=3 per group), and 24 hours later their retinas were harvested and processed for scRNAseq (Fig 1A). At this time point, the bacteria-injected eyes are visibly infected, with pronounced opacity in the anterior chamber (Fig. 1B). Retinal lamination was severely disrupted but not completely destroyed (Fig. 1C). This is ideal as most cells are still alive after the dissociation process (Fig. 1D).

**Figure 1.**
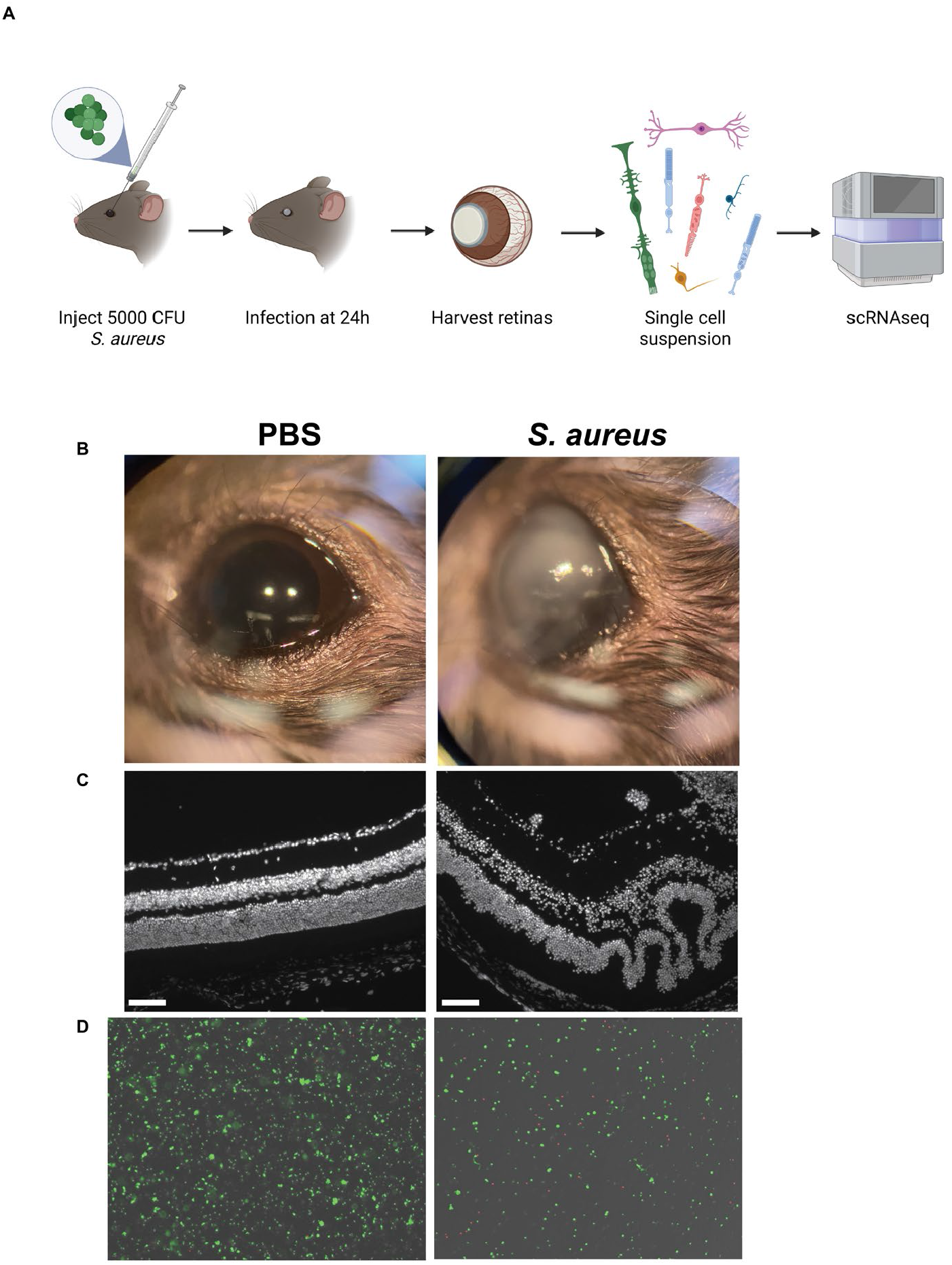
Overview of murine endophthalmitis and scRNAseq. (A) Schematic of protocol.(B) Representative photographs of infected and control eyes 24h post-injection. (C) Representative fluorescent images of retinal cross-sections 24h post-injection. DAPI indicates nuclei. (D) Representative microscopy images of live/dead imaging after retinal dissociation. Green fluorescence: live cells, red fluorescence: dead cells. Scale bars indicate 100 µm.

After quality control of the data, there was a total of 47,011 cells (mean 7,835 cells per retina), and all expected cell types were identified (Fig. 2A). A large population of cells exclusive to the infected retinas were identified as infiltrating neutrophils (Fig. 2B). Automatic cell type labeling was validated by the expression of known markers (Fig. 2C).

**Figure 2.**
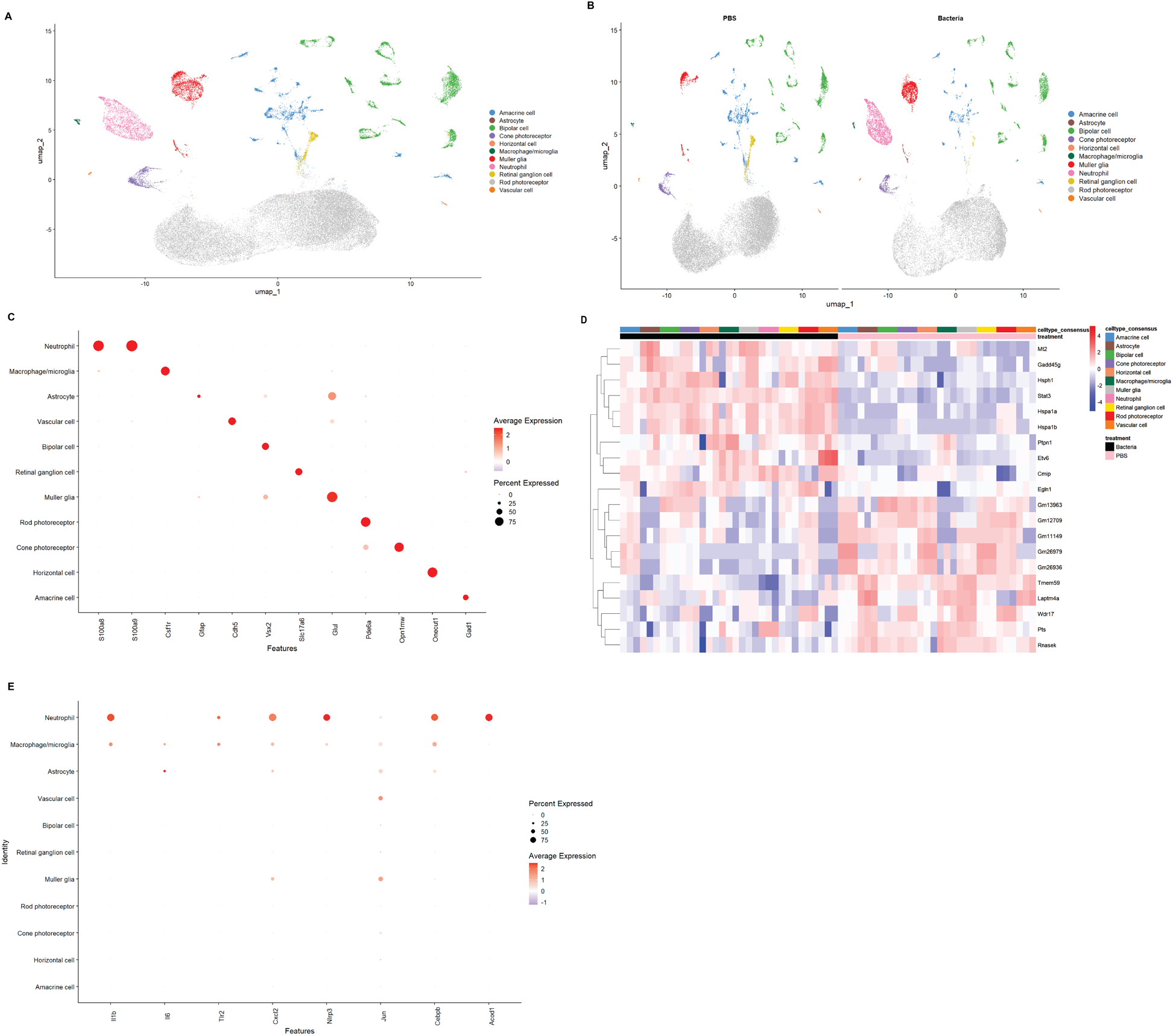
scRNAseq reveals global trends in retinal response to infection. (A) UMAP of cells from all retinas (n=3 PBS, 3 *S. aureus*). (B) UMAP of cells split by treatment group. (C) Bubble plot of cell type markers. (D) Heat map of commonly up- and down-regulated genes across all cell types with infection. (E) Bubble plot of genes previously implicated in murine *S. aureus* endophthalmitis at the bulk-retina level.

Despite the heterogeneity of the retina, some genes changed expression across almost all cell types (Fig. 2D). This included heat-shock proteins (*Hspa1a, Hspa1b, Hsph1*) and the transcription factor *Stat3*, which are known to be upregulated by cellular stress.

Transcriptomic analysis at the bulk-retina level in this model has identified a number of genes significantly upregulated by *S. aureus* infection,^20^ including *Il1b* (pro-inflammatory cytokine), *Nlrp3* (inflammasome mediator), and *Tlr2* (receptor for the *S. aureus* cell wall component peptidoglycan). Neutrophils were the predominant source of these genes, as well as other inflammatory markers (Fig. 2E).

### Evidence of hypoxia in several infected retinal cell populations

Cellular respiration was a top affected pathway among retinal bipolar neurons (Fig. 3A). Examining the genes in this pathway, the majority were significantly downregulated with infection (Fig. 3B). This included genes critical to the mitochondrial respiratory chain, e.g. many in the *Nduf* family. Upregulated genes included the hypoxia markers *Hif1a* and *Bnip3*. Similar trends were also observed in retinal ganglion cells and cone photoreceptors.

**Figure 3.**
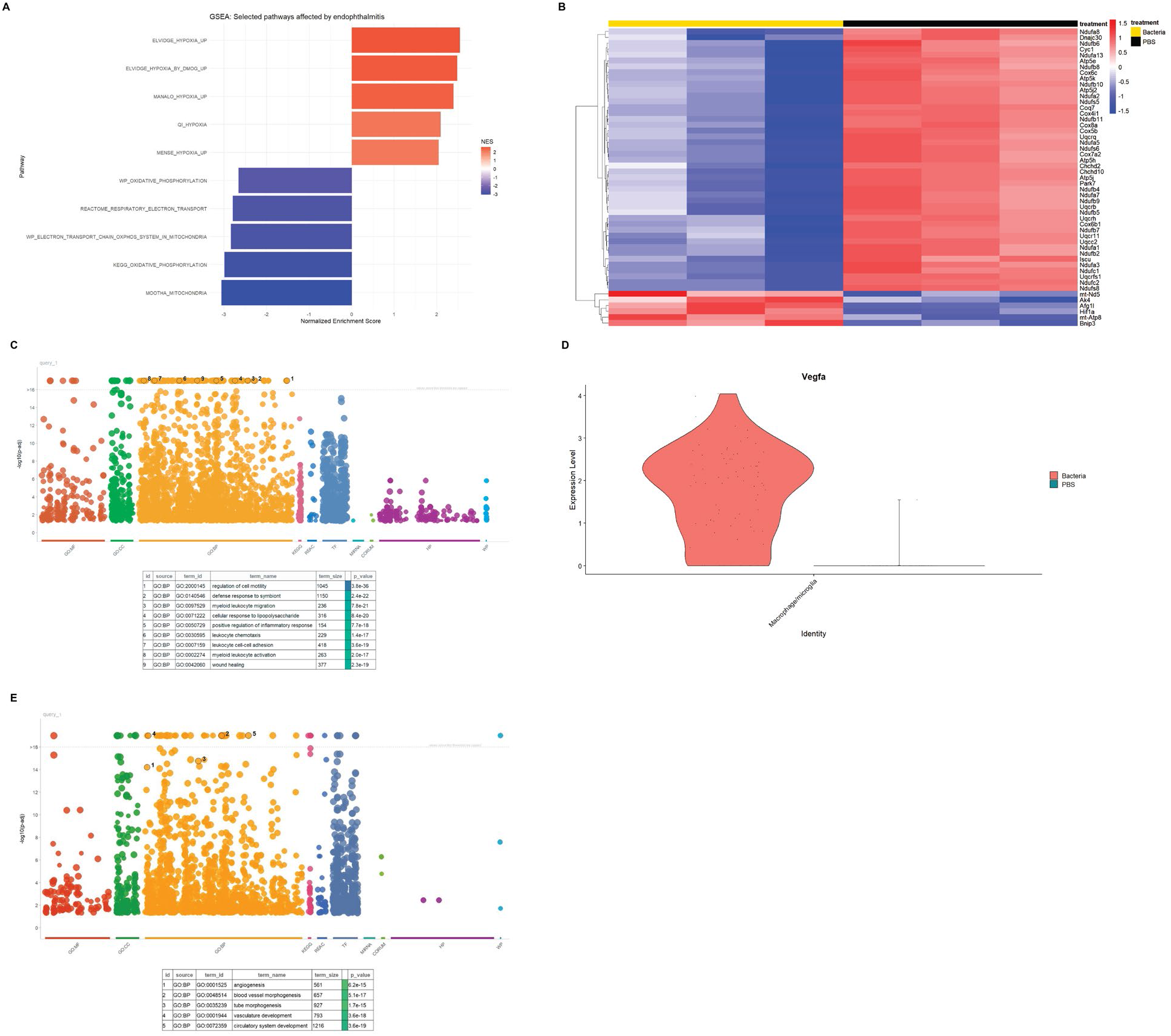
Gene expression changes in macrophage/microglia, vascular cells, and neurons suggest retinal hypoxia. (A) Gene set enrichment analysis in retinal bipolar neurons. (B) Heat map showing expression of cellular respiration genes in retinal bipolar neurons. (C) Gene ontology plot of affected pathways in macrophages/microglia. (D) Violin plot of *Vegfa* expression in macrophages/microglia. (E) Gene ontology plot of affected pathways in vascular cells.

Due to low cell numbers and transcriptional similarity, infiltrating macrophages and resident microglia were combined into a cluster called macrophages/microglia. Endothelial cells and pericytes were combined into a cluster called vascular cells. Gene ontology analysis revealed many pathways affected by infection in monocytes, including wound healing (Fig. 3C). This pathway included significant upregulation of vascular endothelial growth factor A (*Vegfa*) (Fig. 3D).

Vascular cells upregulated angiogenesis markers (Fig. 3E). This does not necessarily mean that neovascularization is occurring in infected retinas, especially since the entire retina may be destroyed first. Instead, this pattern of gene expression is additional evidence for oxygen depletion in the retina.

### Muller glia express ferroptosis markers in response to infection

Expression of *Gpx4*, the main inhibitor of ferroptosis, was strongly decreased in *S. aureus*-infected Muller glia (Fig. 4A). Other markers of ferroptosis were strongly and consistently upregulated in Muller glia: *Acsl4, Hmox1*, and iron importers *Zip14* (*Slc39a14*), *Dmt1* (*Slc11a2*), and *Tfrc* (Fig. 4B). Upregulation of *Slc7a11* would inhibit ferroptosis, but glutathione alone is not sufficient to prevent ferroptosis without Gpx4. Vascular cells also decreased Gpx4, but lacked the other ferroptosis gene expression changes.

**Figure 4.**
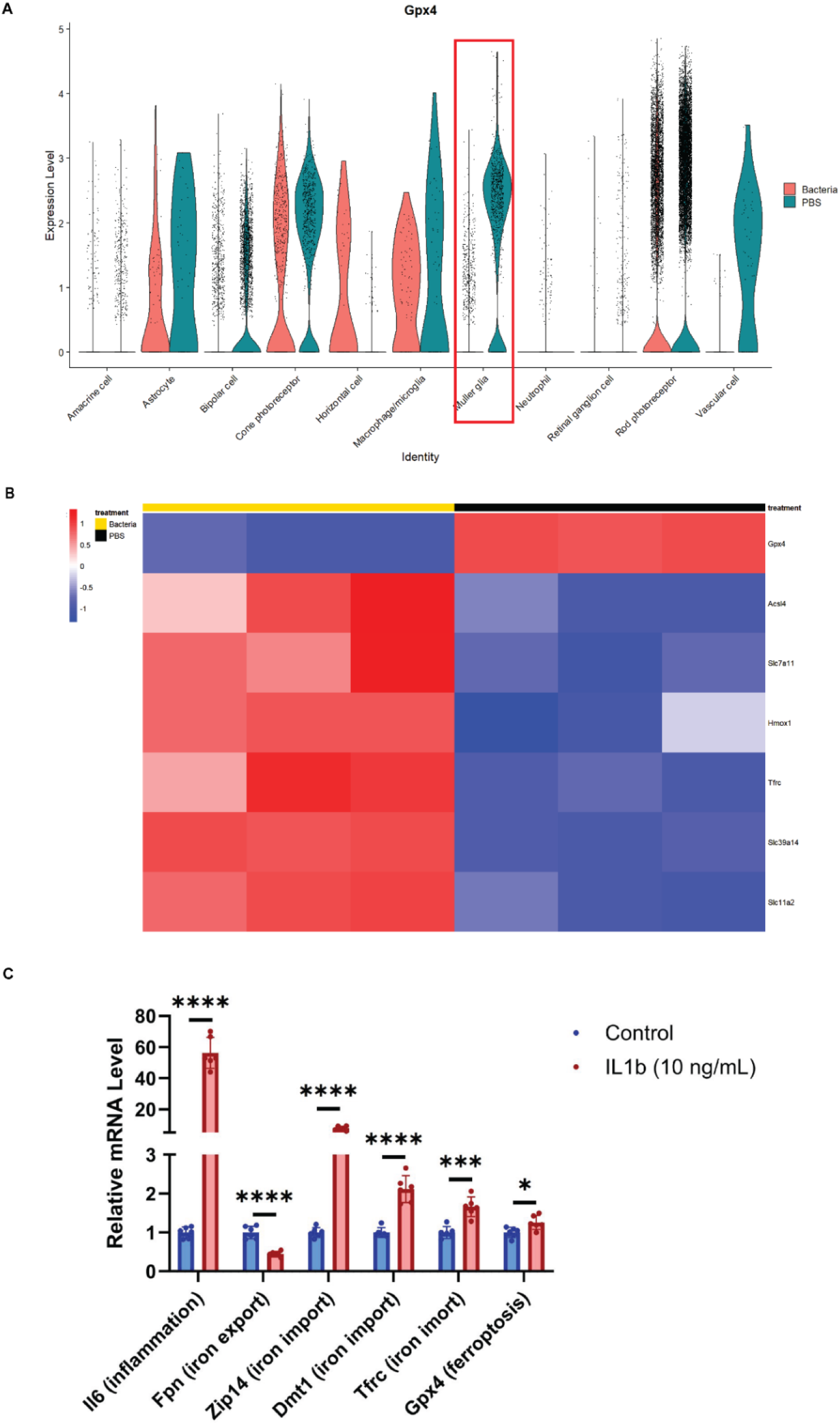
Markers of iron dysregulation and ferroptosis in activated Muller glia. (A) Violin plot of *Gpx4* expression across all cell types. Red box highlights Muller glia. (B) Heat map of iron- and ferroptosis-related genes in Muller glia. (C) qPCR results from MIO-M1 cell culture after 24 hr with or without IL-1β.

To examine this pathway in human cells, the immortalized Muller glia cell line MIO-M1 was used. Cells were incubated with the pro-inflammatory cytokine IL-1β, and expression of ferroptosis-related genes was measured by qPCR (Fig. 4C). IL-1β significantly upregulated *Il6* expression, along with *Zip14, Dmt1*, and *Tfrc. Gpx4* was slightly increased, and *Acsl4* was slightly decreased. The iron exporter *Fpn* also decreased.

## Discussion

*S. aureus* endophthalmitis induces robust retinal inflammation. In our dataset, neutrophils were dominant contributors to the retinal infection response, potentially masking the effect of other cell types when analyzed at the bulk level. The retina may be experiencing hypoxia, affecting both immune and neuronal cells. Muller glia exhibit a pattern of gene expression consistent with iron dysregulation and ferroptosis.

These results confirm a number of pathways and genes known to be altered in this model of endophthalmitis, including inflammation and immune cell infiltration. Bacteria utilize nutrients like oxygen and iron to survive and proliferate, and both hypoxia and iron dysregulation have been associated with infection.^21,22^ However, *S. aureus* can grow in both aerobic and anaerobic environments.^22^ The specificity of the ferroptosis pathway to Muller glia highlights an opportunity for a targeted therapeutic approach.

Retinal damage can continue to progress even once antibiotics have cleared the underlying infection. Heat-killed *S. aureus*, as well as the cell wall component peptidoglycan, can cause retinal inflammation and pathology.^8^ Iron depletion of the vitreous and retinal ferroptosis have been reported with ocular toxoplasmosis infection.^23^ Our data demonstrate that sterile inflammation induces iron dysregulation in human Muller glia in vitro. Overall, this suggests that inflammation alone, rather than pathogen-specific virulence factors, induces cellular iron overload. These data may partly explain why bactericidal treatment alone can be insufficient for resolving endophthalmitis, emphasizing the need for a combinatorial approach.

A critical immune response to infection is the regulation of metal ions. Transition metals like copper, zinc, manganese, and iron are essential for cellular function but cytotoxic at high concentrations, as they can promote the production of reactive oxygen species (copper, iron) or overload the metal-binding domains of enzymes. Macrophages can manipulate the availability of these ions to both starve and overwhelm bacteria, a response called nutritional immunity.^21^

In particular, iron is essential to bacteria (and cellular life in general) as a key enzymatic cofactor. Iron-dependent processes include DNA synthesis, transcription, and cellular respiration.^24,25^ Bacteria upregulate iron-import genes during endophthalmitis infection.^26,27^ This expression pattern occurs across different bacterial species and host organisms, suggesting that iron acquisition is a common objective. Bacteria have evolved effective strategies for surviving in iron-deficient environments.^28^ Iron chelation has shown promise at reducing bacterial load in mouse models of skin wound infection,^29^ sepsis,^30^ and pneumonia.^31^ This includes bacteria which are antibiotic resistant, like MRSA.

Vitreous has an iron concentration of 1 to 3 µM in the human eye.^32,33^ This would be sufficient for *S. aureus* growth, but macrophages can import and sequester iron to dramatically reduce its availability to pathogens.^34^ Therefore, infection leads to a dynamic competition for iron between bacteria and macrophages. The ability of host cells to sequester iron away from pathogens is critical for preventing uncontrolled bacterial proliferation: Knock-out of iron-regulatory proteins (IRPs), from macrophages increases the lethality of *Salmonella* infection in mice.^35^ However, severe or chronic iron accumulation is associated with neuroinflammation and degeneration. In vitro, microglia cultured with iron are more responsive to inflammatory stimuli.^36^ Iron chelation reduces retinal inflammation in non-infectious contexts, like mouse models with features of macular degeneration^37^ and light damage.^38^ Therefore, while inflammation leads to iron import, increased iron in macrophages could also potentiate inflammation. Targeting this pathway could combat the toxic cycle of inflammation while preserving the beneficial immune response.

Müller glia appear to sequester iron as part of their defense against *S. aureus* growth. However, when too much iron accumulates, it can promote the formation of reactive oxygen species, overwhelm antioxidant defenses, and lead to cell death via ferroptosis. Since iron sequestration is an important aspect of immune response, the ideal therapy would prevent ferroptosis while also depriving the pathogen of access to iron.

IL-1β alone was sufficient to induce iron dysregulation, although not ferroptosis, in cultured Muller glia. The likelihood of cell death in vivo may depend on additional factors, including extracellular iron concentration. Muller glia ferroptosis has also been reported in non-infectious models, induced by sodium iodate or silicone oil.^39,40^ These data support the hypothesis that this is a general inflammatory response, rather than a specific response to *S. aureus*.

Endophthalmitis is an aggressive, sight-threatening infection. Even early detection and antibiotic treatment are not always sufficient to prevent irreversible retinal damage. As more intraocular operations and injections are performed every year, and antibiotic resistance spreads among bacteria, the prevalence and seriousness of endophthalmitis is at risk of further increasing. Therapeutics which prevent retinal ferroptosis while sequestering iron from pathogens may have a synergistic effect with antibiotics to combat infection as quickly as possible.

## Notes

### Competing Interest Statement

The authors have declared no competing interest.

### Funding

- National Institutes of Health: R01EY036292, R01EY015240, T32EY007035–46, F31EY035141
- FM Kirby Foundation
- Research to Prevent Blindness
- Paul and Evanina Bell Mackall Foundation Trust

